# What the Phage: A scalable workflow for the identification and analysis of phage sequences

**DOI:** 10.1101/2020.07.24.219899

**Authors:** Mike Marquet, Martin Hölzer, Mathias W. Pletz, Adrian Viehweger, Oliwia Makarewicz, Ralf Ehricht, Christian Brandt

**Affiliations:** Jena University Hospital, Jena, 07747, Germany; Center of Sepsis Control and Care (CSCC), Jena, Germany; RNA Bioinformatics and High-Throughput Analysis, Friedrich Schiller University Jena, Leutragraben 1, 07743 Jena, Germany; MF1 Bioinformatics, Robert Koch Institute, 13353 Berlin, Germany; Institute for Medical Microbiology and Epidemiology of Infectious Diseases, University Hospital Leipzig, Leipzig, 04103, Germany; Leibniz Institute of Photonic Technology (Leibniz-IPHT), Jena, Germany; InfectoGnostics Research Campus, Jena, Germany; Institute of Physical Chemistry, Friedrich-Schiller-University Jena, Jena, Germany

## Abstract

Phages are among the most abundant and diverse biological entities on earth. Phage prediction from sequence data is a crucial first step to understanding their impact on the environment. A variety of bacteriophage prediction tools have been developed over the years. They differ in algorithmic approach, results, and ease of use. We, therefore, developed “What the Phage” (WtP), an easy-to-use and parallel multitool approach for phage prediction combined with an annotation and classification downstream strategy, thus, supporting the user’s decision-making process by summarizing the results of the different prediction tools in charts and tables. WtP is reproducible and scales to thousands of datasets through a workflow manager (Nextflow). WtP is freely available under a GPL-3.0 license (https://github.com/replikation/What_the_Phage).

## Introduction

Bacteriophages (phages) are viruses that infect prokaryotes and replicate by utilizing the host’s metabolism (1,2). They are among the most abundant and diverse organisms on the planet and inhabit almost every environment (2). Phages drive and maintain bacterial diversity by perpetuating the coevolutionary interactions with their bacterial prey, facilitating horizontal gene transfer and nutrient turnover through continuous cycles of predation and coevolution (3,4). They directly impact the microbiome, e.g., the human gut, and can influence human health (5). At the same time, phages in aquatic habitats are responsible for the mortality of nearly 20–40% of prokaryotes every day (6). However, despite having considerable impacts on microbial ecosystems, they remain one of the least understood members of complex communities (7).

The sequencing of the entire DNA of environmental samples (metagenomics) is an essential approach to gain insights into the microbiome and functional properties. It should be noted that due to the genome size of phages between 5 kbp to 500 kbp (8), their entire genome can be sequenced assembly-free via long-read technologies (e.g., Oxford Nanopore Technologies or PacBio) (9). They facilitate phage genome recovery in their natural habitat without the need to culture their hosts to isolate the phages (2) and lead to a rapid increase in human gut virome studies (10). This development demands reliable and easy-to-use phage prediction tools and workflows that can be directly used on assembled sequencing data.

However, predicting phages from metagenomes in general and their differentiation from prophages remains a challenge as there is no established computational gold standard (11). Existing prediction tools rely on direct comparison of sequence similarity (12,13), sequence composition (14,15), and models based on these features derived through learning algorithms (12,13,16,17).

The performance of each prediction method varies (18) depending on the sample type or material, the sequencing technology, and the assembly method, which makes the correct choice for any given sample difficult without having to install and test several tools. To further complicate matters, the user can choose from many tools based on different calculation strategies, software dependencies, and databases. While working with these phage prediction tools, we observed various installation issues and conflicts, making a multi-tool screening approach unnecessary complex, and time-consuming. To overcome these obstacles and issues, we developed “What the Phage” (WtP), a reproducible, accessible, and scalable workflow utilizing the advantages of multiple prediction tools in parallel to detect and annotate phages.

## Design and Implementation

WtP was implemented in Nextflow, a portable, scalable, and parallelizable workflow manager (19). At the time of writing, twelve different approaches to predict phage sequences are included in WtP besides other programs for further annotation and classification. WtP uses so-called containers (Docker or Singularity) for an installation-free workflow execution without dependency or operating system conflicts for each of the currently over 21 programs included. All containers are pre-build, version-controlled, online available at dockerhub.com, and automatically downloaded if used. Additionally, all nine different databases/datasets used by the workflow are automatically managed. The modular code structure and functionalities of Nextflow and Docker/Singularity allow easy integration of other phage prediction tools and additional analysis steps in future releases of the pipeline. The workflow consists of two main phases, which are executed subsequently or, if specified, individually (Figure 1):

**Figure 1:**
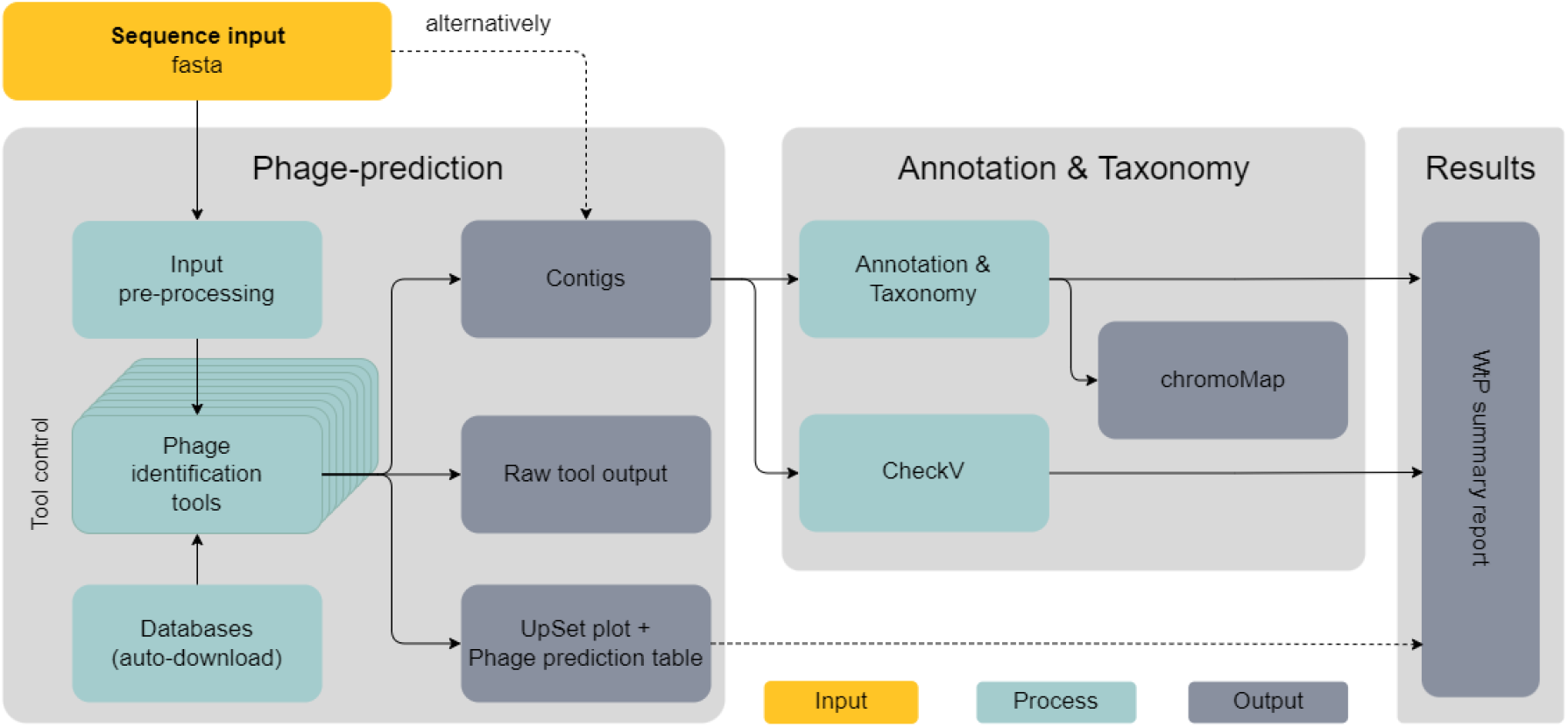
Simplified DAG chart of the “What the Phage” workflow. Sequence input (yellow) can either be first-run through the “prediction” and subsequently “Annotation & Taxonomy” as a whole or used directly as an input for the “Annotation & Taxonomy” only. Each of the multiple phage prediction tools can be individually controlled if needed (tool control).

1. Prediction: The prediction of putative phage sequences
2. Annotation & Taxonomy: The gene annotation and taxonomic classification of phage sequences

### Prediction and Visualization

The first stage takes a multi-fasta file as input (e.g., a metagenome assembly), formats it to the demands of each tool, and filters sequences below a user-defined length threshold (1,500 bp by default) via SeqKit v0.10.1 (20). Sequences that are too small usually generate false-positive hits, as Gregory *et al*. (21) observed. The phage prediction process is performed by eleven different tools in parallel: VirFinder v1.1 (15), PPR-Meta v1.1 (17), VirSorter v1.0.6 (with and without virome mode) (13), DeepVirFinder v1.0 (22), Metaphinder with no release version (using default database and own database (Zheng *et al*.)) (23), Sourmash v2.0.1 (14), Vibrant v1.2.1 (with and without virome mode) (12), VirNet v0.1 (24) Phigaro v2.2.6 (25), Virsorter2 v2.0 and Seeker (26) with no release version. Tool outputs are collected in a detailed result report (See section: Result report, Figure 2, https://replikation.github.io/What_the_Phage/).

**Figure 2:**
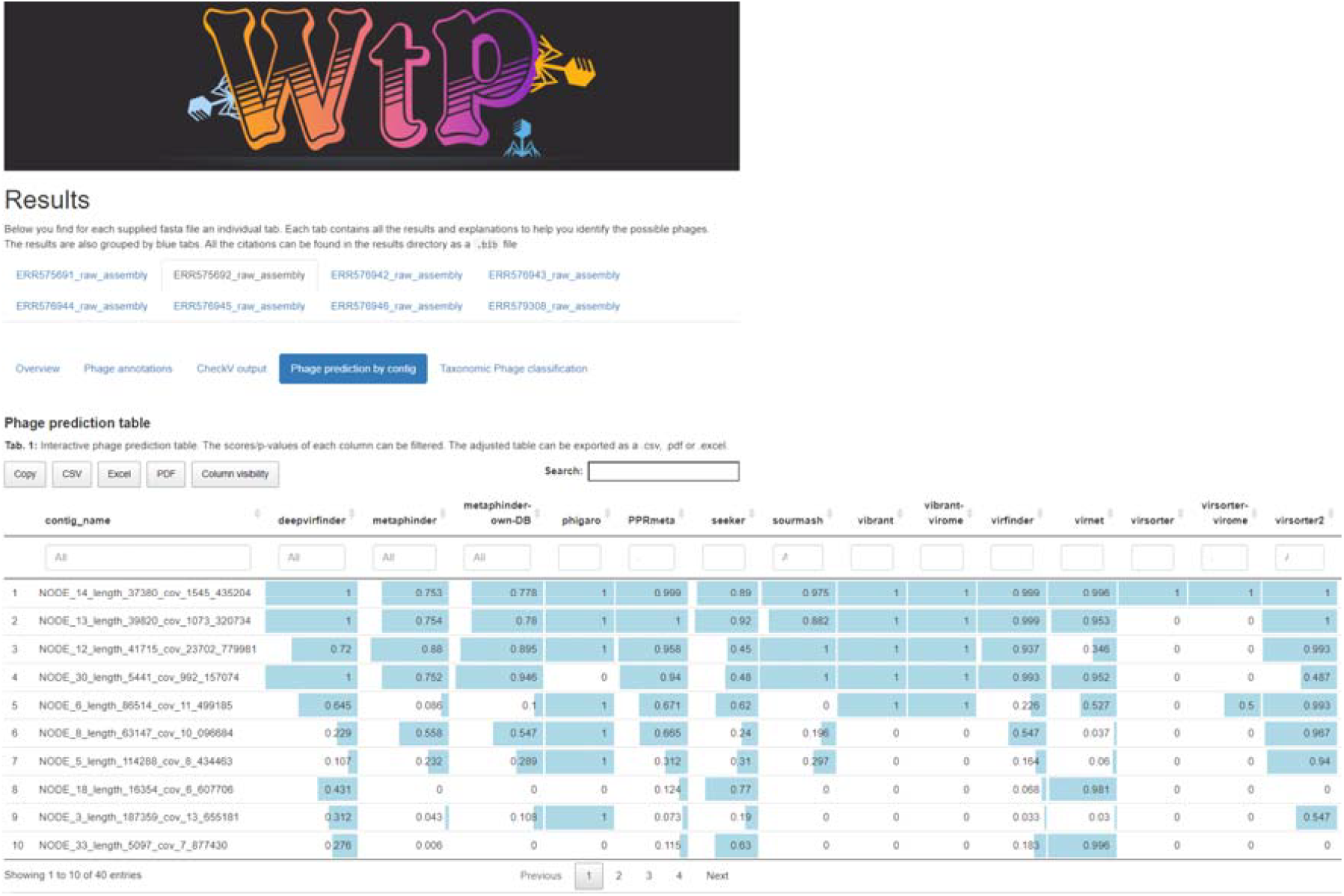
*Example of the final report showing* the analyzed sample ERR575692 *with the “Phage prediction by contig table” section opened*.

### Functional annotation & Taxonomy

For this step, phage positive contigs are used and either automatically retrieved from the prediction step or directly via user input. Prodigal v2.6.3-1 (27) is used in metagenome mode to predict ORFs and HMMER v3.3 (cutoff: -E 1e-30) (28) to identify homologs via the pVOG-database (29). All annotations are summarized in an interactive HTML file via chromoMap (30) (see Figure 3). Additionally, WtP classifies all contigs via sourmash and provides a probability score to the corresponding taxonomic classification based on Zheng *et al*. database.

**Figure 3:**
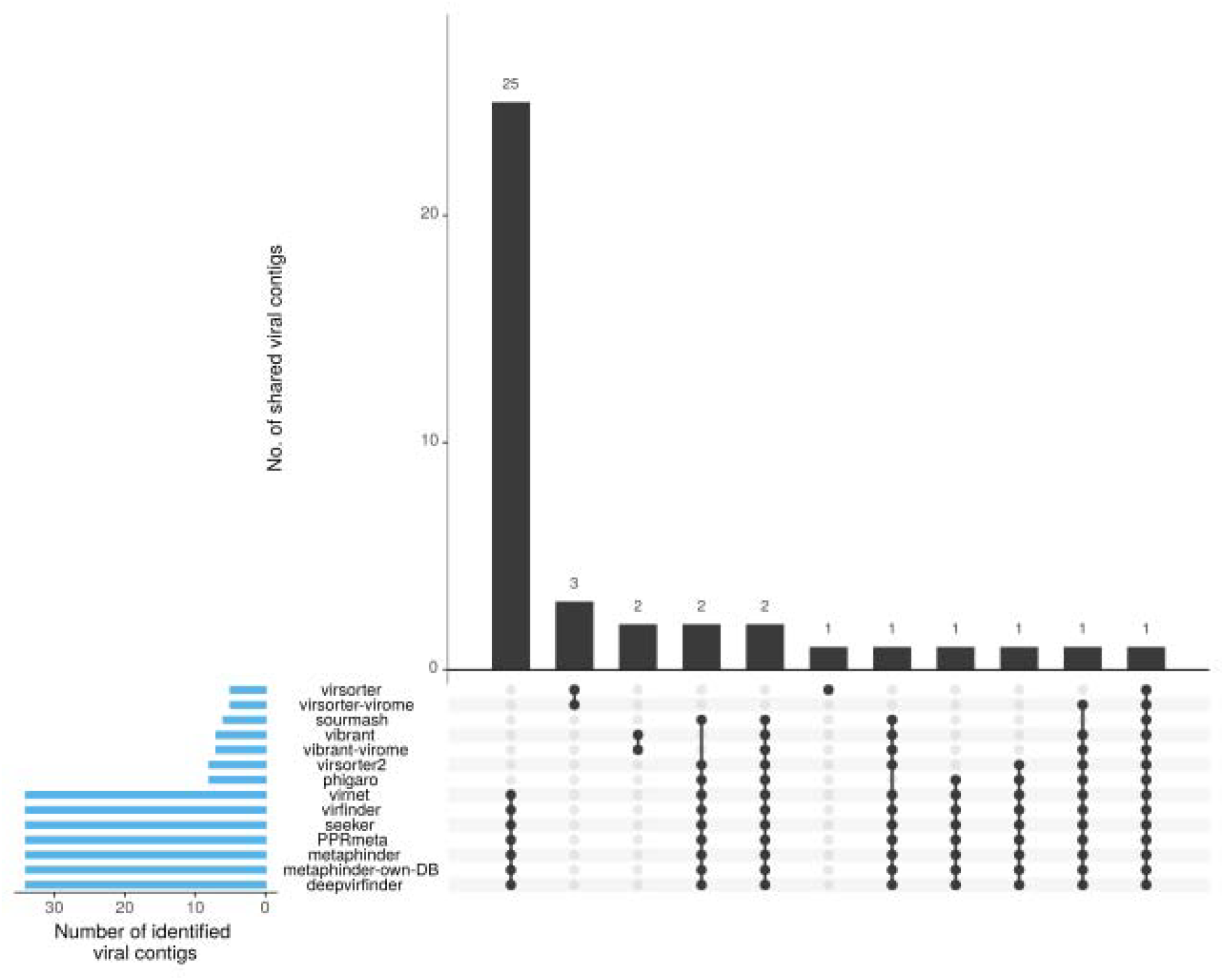
UpSet plot summarizing the prediction performance of each tool for the sample ERR575692. The total amount of identified phage-contigs per tool is shown in blue bars on the left. Black bars visualize the number of contigs that each tool or tool combination has uniquely identified. Each tool combination is shown below the barplot as a dot matrix.

### Result report

WtP streamlines the detection of phage sequences across multiple tools in their default settings, thus balancing some drawbacks (e.g., relying on updated databases, only predicting phages available in databases). To ease the data interpretation for the user, WtP collects the results in a detailed summary report HTML file for simplified interpretation (Figure 2, full report on: https://replikation.github.io/What_the_Phage/). The report contains an UpSet plot summarizing the prediction performance of each tool (Figure 2). Finally, the “phage prediction by contig table” (Figure 2) summarizes tool outputs for each contig based on the tool’s output. WtP assigns numeric values to tools that do not generate p-values or scores between 0 and 1 (see result report, Phage prediction by contig section) and sorts them based on phage likelihood. All the results are individually filterable so the user can consider additional insights or information provided by community platforms such as IMG/VR, iVirus, or VERVE-NET.

### Other features

All mandatory databases and containers are automatically downloaded when the workflow is started and stored for the following executions. Additionally, the workflow can be pre-setup to analyze sequences offline subsequently. WTP provides the raw output of each tool to support a transparent and reproducible mode of operation. Maximum execution stability is ensured by automatically excluding phage prediction tools that cannot analyze the input data without failing the workflow (e.g., file too large, not the scope of an individual tool). We also provide a detailed user manual that is regularly updated and available at www.mult1fractal.github.io/wtp-documentation/.

### Dependencies and version control

WtP requires only the workflow management software Nextflow (19) and either Docker or Singularity (31) installed and configured on the system. The pipeline was tested on Ubuntu 16.04 LTS, Ubuntu 18.04 LTS, and Windows 10 (via Windows Subsystem for Linux 2 using Docker). The installation process is described in detail at mult1fractal.github.io/wtp-documentation/. Each workflow release specifies the Nextflow version the code was tested on to avoid any version conflicts between the workflow code and the workflow manager at any time. A specific Nextflow version can be directly downloaded as an executable file from https://github.com/nextflow-io/nextflow/releases. Additionally, each container used in the workflow is tagged by the accompanying tool version, pre-build, and stored on hub.docker.com.

## Results

To demonstrate the utility and performance of WtP, we analyzed a described metagenome data set (ENA Study PRJEB6941, ERR575692) using a local desktop machine (24 threads, 60 GB RAM, Ubuntu 18.04.4 LTS) and WtP release v1.1.0. In this study (32), Kleiner *et al*. sequenced an artificial microbiome sample which was produced via bacteria and phage cultures in mice feces (germ-free C57BL/6 J mice). The group added six different phages: P22, T3, T7, □6, M13, and □VPE25 and two bacteria (*Listeria monocytogenes and Bacteroides thetaiotaomicron*) to germ-free feces. We, therefore, expect the prediction of the six known phages and possibly other phage sequences related to both bacteria strains. Still, false-positive hits and tool disagreements are plausible results during the phage prediction step.

The raw read data set composed of eight samples was downloaded from the ENA server and individually assembled via metaSPAdes v3.14 using the default settings (33). The resulting assembly files (available at https://github.com/mult1fractal/WtP_test-data/tree/master/01.Phage_assemblies) were analyzed with WtP (release v1.1.0, default settings). As WtP uses multiple tools for phage prediction, an UpSet plot summarizes for each sample the performance of all approaches executed successfully (Figure 3 for sample ERR575692).

In general, the prediction values are high (>0.7) for the first four sequences (NODE_14, NODE_13, NODE_12, NODE_30), indicating high consensus among the prediction tools, although in some cases tools prediction values were below 0.5 (Phigaro: NODE_30, Seeker: NODE_12 and NODE_30, Virnet: NODE_12 and Virsorter2: NODE_30) (Supplementary Figure S1).Prediction values for NODE_6 are below 0.67, and Virsorter2 and Phigaro show high values > 0.99. The same applies to NODE_8 and NODE_5, indicating dissonance for these three contigs. Surprisingly, Virsorter and Virsorter-virome only predict contig NODE_14 as phage. In case of dissonance and when tools coincide, validation of contigs via phage annotations and CheckV simplifies the assessment of each contig as a phage or not. In our case, phage genes (like tail and capsid genes) were annotated on all seven contigs (Figure 4).

**Figure 4:**
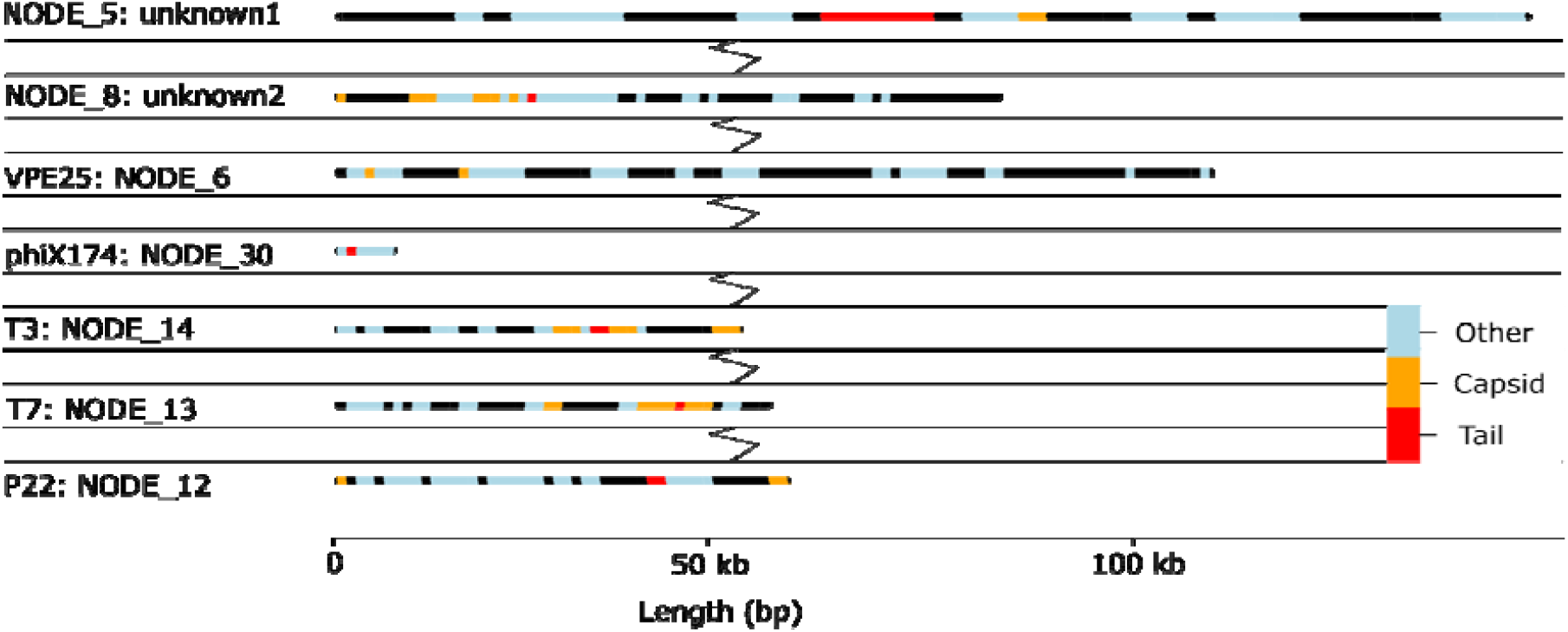
Visual annotation of phage contigs and annotated protein-coding genes via chromoMap. Annotations are colored based on the categories of capsid genes (orange), tail genes (red), and other genes (blue). For better readability, other contigs without either capsid or tail genes have been removed from this figure.

The workflow was able to detect contigs that correspond to the phages P22 (NODE_12), T3 (NODE_14), T7 (NODE_13). In addition, the phage for the internal Illumina control (phiX174: NODE_30) was also identified. The M13 phage (27) could not be identified as it was not assembled via metaSPAdes due to the low read-abundance and low coverages (below 0.55x, determined by Kleiner *et al*. (27)). The same applies to phage □6, which was not detectable by Kleiner *et al*. (27). However, VPE25 (NODE_6) was initially not taxonomically classified by WtP as it was not represented in the taxonomic database (Zheng *et al*.) at this time; however, the corresponding contig was annotated with essential phage genes (Figure 4). Therefore, this unclassified contig was compared manually via blastn (nr/nt database) and resulted in the genome sequence of VPE25 (PRJEB13004).

Furthermore, CheckV determined a phage completeness score of over 89% for all seven contigs (Table 1). In addition to the phages mentioned above, two more large contigs with capsid and tail gene annotations indicate prophage(s) of *Salmonella enterica* (contig NODE_5 and NODE_8). Both contigs showed tail and capsid genes and were labeled as prophages via CheckV with estimated completeness of over 99.99 %. These results were manually confirmed using NCBI’s blastn (nr/nt database). The sequences matched with 100% identity to *Salmonella enterica (Salmonella enterica strain FDAARGOS_768 chromosome, complete genome)*, but not to prophage sequences.

**Table 1:** Summary of the CheckV output for the sample ERR575692. All contigs with a completeness > 89 % and a length > 5,000 bp are shown.

| Phage name | Contig_id | Gene count | CheckV quality | Completeness | contig length [bp] |
| --- | --- | --- | --- | --- | --- |
| unknown1 | NODE_5 | 107 | Complete | 100.0 | 114,288 |
| unknown2 | NODE_8 | 71 | High-quality | 100.0 | 63,147 |
| VPE25 | NODE_6 | 137 | High-quality | 99.99 | 86,514 |
| phiX174 | NODE_30 | 8 | Medium-quality | 89.35 | 5,441 |
| T3 | NODE_14 | 43 | High-quality | 93.34 | 37,380 |
| T7 | NODE_13 | 53 | Complete | 99.48 | 39,820 |
| P22 | NODE_12 | 67 | Complete | 100.0 | 41,715 |

Some limitations must be noted. No specialized phage assembly strategy or any cleanup step was included during the assembly step. Therefore, some smaller mice host contigs (below 5,000 bp) produced false positive hits. However, these contigs were clearly distinguishable after the “Annotation & Taxonomy” step both in CheckV and due to the lack of typical genes related to, e.g., capsid or tail proteins, showing the application of WtP also for contaminated datasets. WtP does not filter the output of phage prediction tools for prophages, although the CheckV output indicates if a contig could be a prophage.

## Conclusion

With the rise of metagenomics and the application of machine learning principles for virus detection, several phage prediction tools have been released over the last few years. All these tools utilize a variety of prediction approaches, all with advantages and limitations. The user’s choice for using certain tools often depends strongly on its usability and accessibility and less on its performance. While some tools already come with a packaging system such as Conda or a containerized environment, there exists no general framework for their execution database dependencies, and installation issues prevent many potential users from using certain tools. At least one multitool approach was implemented on a smaller scale by Ann C. Gregory *et al*. (comprising only VirFinder and VirSorter) (20).

The overarching goal of WtP is to make phage prediction tools more accessible for a broader user spectrum and non-bioinformaticians, as culture-free sequencing led to the rapid increase of phage studies (10). WtP acts as an ideal all-encompassing starting point for any given assembly while providing a searchable and filterable report of the analyzed data. The user is provided with sufficient processed data (such as tool performance comparisons, taxonomic assessments, and annotation maps) to work reliably with the predicted sequences and support the decision-making process if different prediction tools are not in agreement with each other. For this, further information and guides are provided either via the report or the hosted manual. WtP streamlines the prediction of phage sequence recognition across multiple tools in a reproducible and scalable workflow to allow researchers to concentrate on their scientific questions instead of software implementations.

### Future directions

WtP is a workflow project that will be improved and extended as the modular approach and containerization simplify the integration of new tools. The predictive scope of WtP can be extended to other viruses (such as RNA viruses) and prophages by including future tools specifically designed for such use cases and by adjusting filter and annotation steps. Furthermore, we plan to support the input of long raw reads as an alternative to assemblies. The versioning of WtP represents a well-functioning approach with tested and up-to-date versions of the workflow. Thus, the correct functioning of the workflow is always guaranteed and allows a reliable and fast prediction of phage sequences.

## Declarations

### Availability

Source code: https://github.com/replikation/What_the_Phage

Result Report: https://replikation.github.io/What_the_Phage/

WtP result data storage: https://osf.io/kuc96/

WtP databases: https://osf.io/wtfrc/

Sequence data used in this work is available at: https://github.com/mult1fractal/WtP_test-data

## Acknowledgments

We thank Michael Shamash for his help in properly testing and validating WtP on a Slurm-based HPC utilizing Singularity, Luiz Irber, to improve the sourmash integration. We also thank Polina Tikhonova and Nikos P. for their help in implementing their phage prediction tools Phigaro and Seeker.

